# Absence of general rules governing molluscan body-size response to climatic fluctuation during Cenozoic

**DOI:** 10.1101/438002

**Authors:** Debarati Chattopadhyay, Devapriya Chattopadhyay

## Abstract

Body size is a key factor in dictating the fate of interaction between an organism and its surrounding environment. A negative temperature-size relationship (TSR) has been suggested as one of the universal responses to climatic warming. It is also predicted that groups with narrow latitudinal range, tropical affinity and higher body size, would show higher sensitivity to climatic fluctuation. Moreover, because of the difference in thermal sensitivity, it is also expected that the response to climatic fluctuation would be different between epifaunal and infaunal groups. To confirm the generality of these relationship among marine families, we compiled the relationship between body-size and global temperature trends over Cenozoic using a database of marine benthic molluscs of class gastropoda and bivalvia resolved to temporal stages. We evaluated the dependence of climate induced body-size response to the existing size and latitudinal spread via correlating the first-difference correlation coefficient of temperature-size (ρ^1st^ _(size-temp)_) with maximum size and latitudinal spread of family respectively. Cenozoic record of this highly diverse group does not show any signature of TSR for molluscan class or for any other regional, ecological groups during the past 66My long climatic fluctuation. We did not find any evidence supporting heightened response to climatic fluctuation in groups with limited latitudinal spread or with large body-size. The tropical species did not show significant difference in their body-size response in comparison to temperate species. It also shows lack of any difference in response between ecological groups of molluscs with varying substrate relationship and hence, refutes the predicted variation due to difference in thermal specialization. Although a negative correlation between maximum latitudinal spread and ρ^1st^ _(size-temp)_ is observed for infaunal families, it is not statistically significant. Our results highlight the limited validity of “universal rules” in explaining the climate induced morphological response of marine communities in deep time and underscores the complexity in generalizing the biotic outcome of future climatic fluctuation.

## Introduction

Body size of an organism has long been considered as one of the most important biological parameters because it is related to various physiological, life-history and ecological traits influencing community structure and functioning of ecosystem [1–7]. Variations in body size, although studied for many terrestrial ectotherms and endotherms, remains poorly understood for marine ectotherms especially in the context of temperature influence [8–11]. Changes in body size have important implications for the thermal biology and energetics of endotherms and ectotherms because body size directly affects energy and water requirements for thermoregulation, energy and life-history characteristics [2, 12, 13]. Change in body size will, therefore, have consequences for resilience of a group in the event of climate change. It has been recognized that many of the biological predictions about body size response to climatic variation, that have been developed primarily based on data from terrestrial vertebrates, are nonexistent or lack significance when applied for marine organisms [10]. Limitation of such extrapolation regained considerable interest in the present decades, primarily because of its relevance in the biotic response to recent climate crisis of global warming. The overall effect of global warming on ecosystem is appreciated, but general rules to predict the fate of a specific group, as it responds to climate warming, is largely absent. This is especially true for marine ectotherms, though they represent more than 80% of the marine species on Earth [14, 15].

A negative relationship between temperature and body size, known as temperature-size rule (TSR) considered as one of the few general rules and is supported by many recent and past marine records [10, 11, 16–18], but counterexamples also exist [19]. Moreover, studies supporting the existence of TSR have been primarily performed at population and community level; it is yet to be validated at the species level using wide temporal data. It is important to note that the response of body size to climatic shift might not be homogeneous across groups [20] and individual ecology may play a significant role in guiding the response. It has been speculated that climate warming is expected to affect body size of groups adapted to various climate with different severity. For example, species that are restricted to a narrow latitudinal range, such as stenotherms and tropical species, would show higher degree of body size change during any climatic fluctuation compared to widespread taxa [14, 20]. Temperate species occupying wider ranges of habitats, hence phenotypically more variable than tropical species, should respond differently [21]. On a similar note, body size change is thought to be dependent on existing body size of a species due to size specific difference in thermal sensitivity [22, 23] and hence, smaller individuals are more likely to be affected by climatic fluctuation. Moreover, it has been recognized that there is a strong ecological selectivity in response of the biota to climatic shifts [24]. It has been observed that short-term exposure to high or low temperature usually causes increased tolerance to acute thermal extremes whereas long-term exposure to moderate temperatures can induce heightened thermal sensitivity [25, 26]. Consequently, we expect a difference in the thermal specialization and consequent body size response between epifauna and infauna.

It is well established that climate induced variability in body size could take decades to become observable and hence can easily be missed in ecological timescale [8, 18, 23, 17]. Because the effect of climatic fluctuation on body-size is difficult to model from a single event of present global warming or to simulate from laboratory experiments due to its complexity [6], ancient climatic fluctuations and associated body-size response provide critical insight. One way of addressing this issue would be to look at the fossil record of marine groups and evaluate the pattern of body size change during geologic intervals with marked climatic shifts. Molluscan group fits the description perfectly because they have high preservation potential and experiences different levels of temperature fluctuation depending on their lifemode [28–30]. Cenozoic marks a time period with numerous paleoclimatic fluctuations that have been reconstructed with well constrained data [31]; moreover, Cenozoic fossil record is more complete [32] compared to the earlier periods. Therefore, Cenozoic molluscan fossil record gives the ideal scenario to test the following hypotheses:

i. The existence of a negative TSR for molluscan body size during last 66Ma.
ii. Families of smaller body size and narrow latitudinal range should show higher magnitude of change.
iii. The body-size response of species with tropical affinity should be different in comparison to temperate species.
iv. The body-size response of species that are infaunal should be much more pronounced than epifaunal species.

In this article, we tested these hypotheses using data on body size and ecological information for 10,388 occurrences of marine benthic molluscs of class gastropoda and bivalvia from the Paleobiology Database (paleodb.org), with published paleotemperature estimates [33] resolved to temporal stages.

## Materials and methods

### Dataset compilation

Two classes of benthic molluscs, bivalvia and gastropoda, from 18 established stages of Cenozoic are used for this study. The other classes of molluscs either have a poor fossil record or are not contributing significantly during Cenozoic [34]. Our database contains 10,388 fossil occurrences representing 154 unique families (gastropod-97, bivalve-57). We traced the temperature change over last 66 my using the curve published by Zachos et al. [33] based on stable oxygen isotopic variation during Cenozoic. Using the variation in temperature within each of the 18 time bins, we calculated stage-specific mean temperature.

Data on body-size, ecological character, stratigraphic range and latitudinal distribution for all the species were collected from the Paleobiology Database by downloading “measurements of specimens” for specific geologic stages (http://fossilworks.org/; date: 17^th^ March, 2016). Each downloaded datasheet for a specific time bin contained the occurrence data of species along with location, size and number of specimens used for measurements. A total of 15,554 individuals were used for the size measurements (S1 Table). We considered the greatest dimension (length or width) as a proxy of size. For each species, we extracted size information along with the specific latitude where they have been recorded in a particular time bin. This size is the geometric mean of measurements of multiple specimens belonging to a single species in a specific time bin. The size data is then log transformed (base 10) for all subsequent analyses. Hence, a particular species is represented by multiple size information in a single time bin if the species appeared in multiple locations. A specific time bin, therefore, contains body-size of various species belonging to different families.

For calculating ρ^1st^ _(size-temp)_ for a specific family, we considered maximum body-size of a species (belonging to that specific family) in a specific time bin. Hence, various time bins can be represented by different species of the same family. These family specific ρ^1st^ _(size-temp)_ is later used for evaluating its dependence on body size and latitudinal spread. For Fig 3, the x-axis value (Maximum size) of any point represents the largest measures of the body-sizes of all species belonging to a particular family over Cenozoic. For Fig 4, the same method is used to calculate maximum latitudinal spread from the tropics of a family over Cenozoic. This exercise was conducted for all families and then dividing them into ecological groups (infauna, epifauna).

We assigned each species to an ecological group based on their substrate relationship. We divided the species into two groups (tropical and temperate) based on their latitude of occurrence. A specific species is considered to be a tropical species if its occurrences during Cenozoic is restricted between 23.5° N to 23.5° S. A similar rationale was followed to identify temperate species when a species occurrence is strictly restricted between 23.5° to 66.5° in each hemisphere throughout Cenozoic. If a species appears in both tropical and temperate latitudes (mixed latitudinal zone), they are not considered in the analyses.

### Statistical analyses

In order to evaluate the general trend of body-size response during climatic fluctuation, we correlated individual size measures (average, minimum and maximum) with temperature binned over Cenozoic. We used correlation test on values detrended by a first-differences transformation of species level data [35].

In this correlation test, we excluded a total of 69 families from our dataset because they are either present in less than 4 time bins or the size variation between any two consecutive bins is zero. Rest of the 85 families (gastropod-54, bivalve-31) are later used for testing the dependence of families on existing body-size and latitudinal spread. Mean body-size, although, gives the central tendency of overall data [36], the effect of any temperature fluctuation is more likely to be observed in maximum body-size [37]. Hence, we used the correlation between maximum body-size of each family and their corresponding ρ^1st^ _(size-temp)_ to evaluate the effect of existing body-size of families (“existing size dependence”). We used the correlation between maximum latitudinal spread of individual family and their corresponding ρ^1st^ _(size-temp)_ to evaluate the effect of latitudinal extent of a family on its body-size response to climatic fluctuation (“latitudinal spread dependence”). The same exercise is done on a subset of the data based on their ecological character. We used Pearson’s correlation test to evaluate the direction and magnitude of all relationships for this study. Calculating significance for multiple comparison, we used False discovery rate procedure described in Curran-Everett, [38]. To evaluate the difference in body size response between pairs (infauna-epifauna, tropical-temperate), we used Cramer test. All the analyses are performed in R software [39].

## Results

The congruence between body size and temperature is observable only for raw data (ρ= −0.01; P= 0.02) (Fig 1) when body size and temperature estimates are both binned by geological stages; however, this relationship is not found in the detrended data, corrected for autocorrelation by first differencing (ρ= 0.18; P= 0.48) (Fig 2A), implying a lack of size-reduction due to climatic warming during Cenozoic. This lack of temperature-size relationship holds true in a wide range of data treatments, such as considering different measures of size (maximum, minimum) (Fig 2B-C, Table 1).

**Fig 1.**
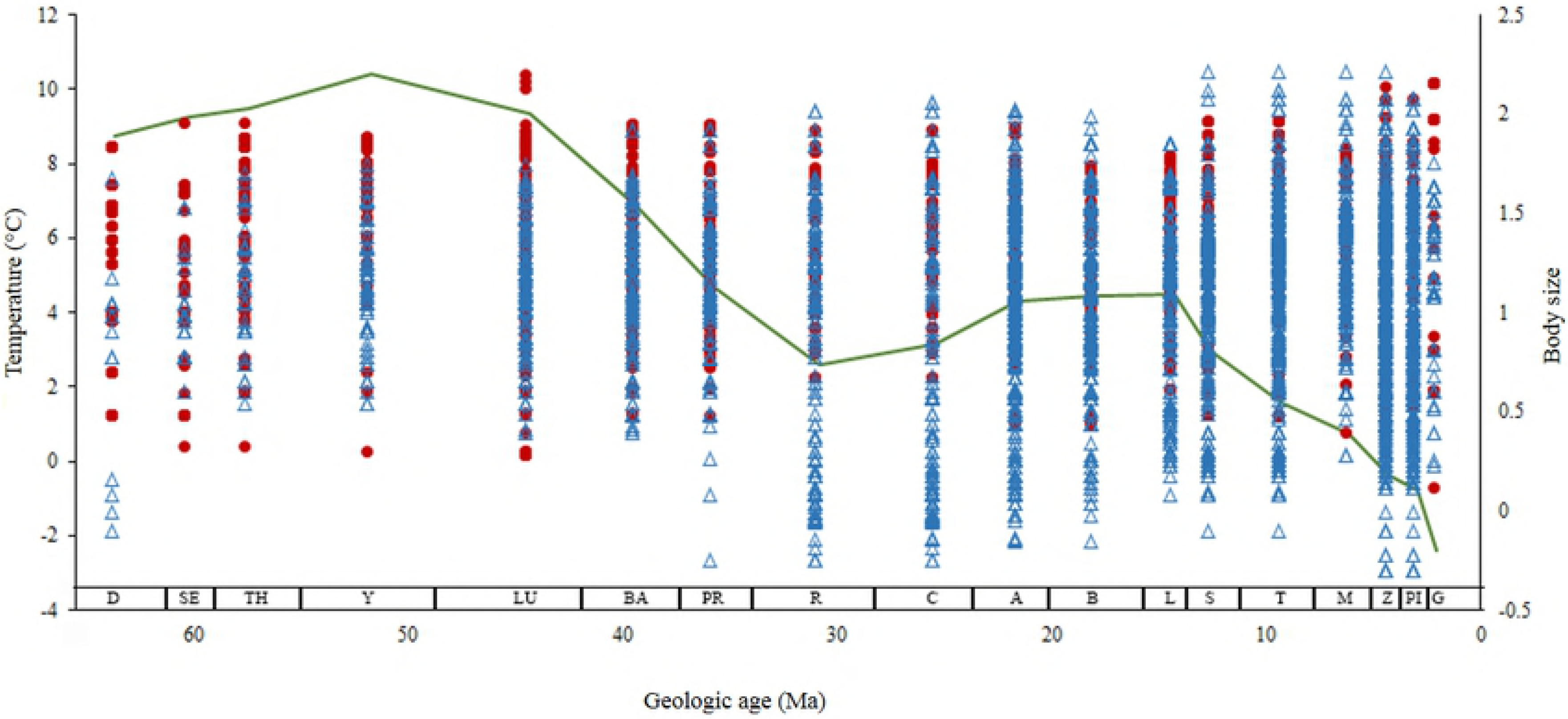
Cenozoic history of climatic fluctuation and body-size change of marine benthic molluscs. The relationship between mean body-size and mean temperature (green line) is showing a negative relationship. The red circles represent bivalves and the blue triangles represent gastropod species occurrences. Size data is log transformed. G, Gelasian; PI, Piacenzian; Z, Zanclean; M, Messinian; T, Tortonian; S, Serravallian; L, Langhian; B, Burdigalian; A, Aquitanian; C, Chattian; R, Rupelian; PR, Priabonian; BA, Bartonian; LU, Lutetian; Y, Ypresian; TH, Thanetian; SE, Selandian; D, Danian.

**Fig 2.**
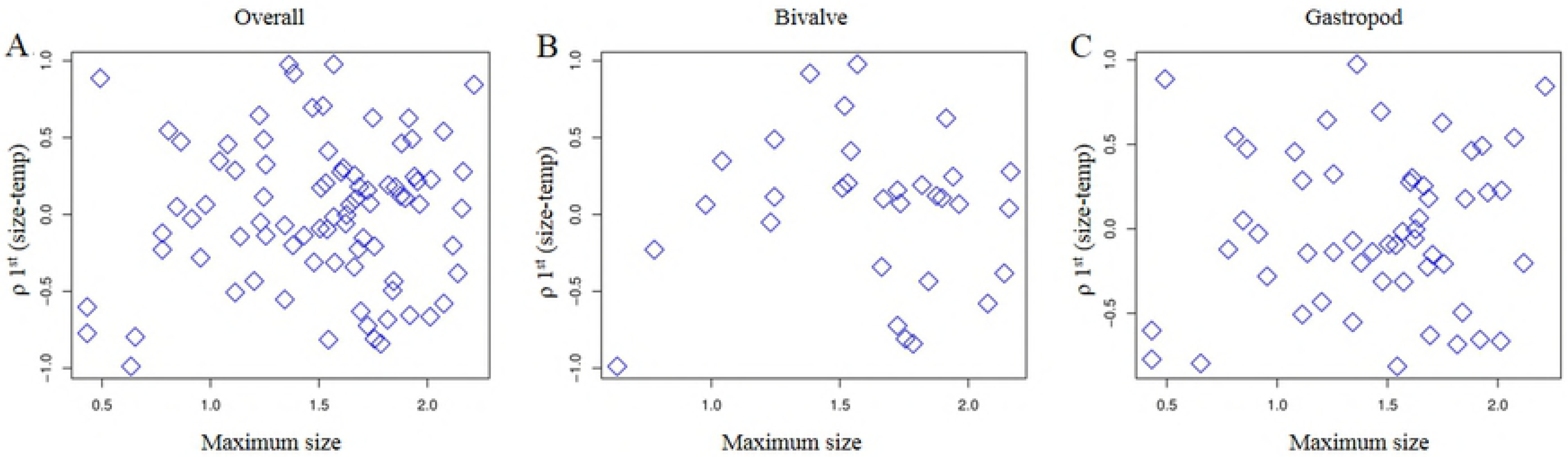
Detrended data (first difference values binned by geologic stages) is not showing any evidence of climate induced body-size reduction in last 66Ma. The detrended relationship between body-size (blue) and temperature (orange) considering average body-size (A), maximum body size (B) and minimum body-size (C). Size data is log transformed.

**Table 1.** Relationship between body-size and temperature during Cenozoic.

|  |  | Mean |  | Maximum |  | Minimum |  |
| --- | --- | --- | --- | --- | --- | --- | --- |
| | | $\rho^{1st}_{Temp}$ | P | $\rho^{1st}_{(Size-Temp)}$ | P | $\rho^{1st}_{(Size-Temp)}$ | P |
| Overall |  | 0.18 | 0.48 | -0.11 | 0.69 | 0.09 | 0.73 |
| Latitudinal zone | Tropical | 0.07 | 0.8 | 0.42 | 0.09 | 0.005 | 0.98 |
|  | Temperate | 0.23 | 0.36 | -0.12 | 0.66 | 0.23 | 0.37 |
| Substrate relationship | Epifauna | 0.1 | 0.7 | - 0.11 | 0.67 | 0.12 | 0.64 |
|  | Infauna | 0.29 | 0.3 | 0.08 | 0.77 | -0.2 | 0.44 |

The significance is set at a value determined by False discovery rate procedure.

No correlation is observed between maximum body-size and ρ^1st^ _(size-temp)_ i.e. the coefficient of correlation between 1^st^ difference of maximum body-size vs temperature for overall data, bivalves or gastropods (Fig 3). Same is true for latitudinal spread and ρ1st (size-temp) (Fig 4) (Table 2)

**Fig 3.**
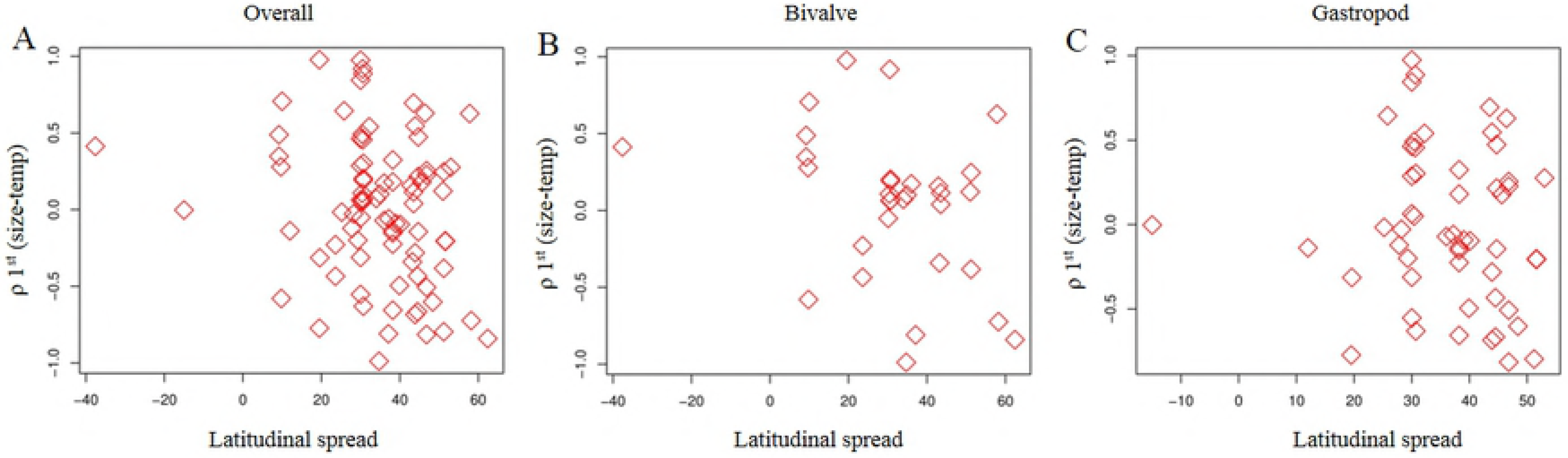
Climate induced body-size response has not been affected by the existing size. Dependence of climate induced body-size response on existing body-size of families of all (A), bivalves (B) or gastropods (C) families during Cenozoic Era. The vertical axis represents the coefficient associated with detrended correlation between temperature and mean body-size at family level. Size data is log transformed.

**Fig 4.**
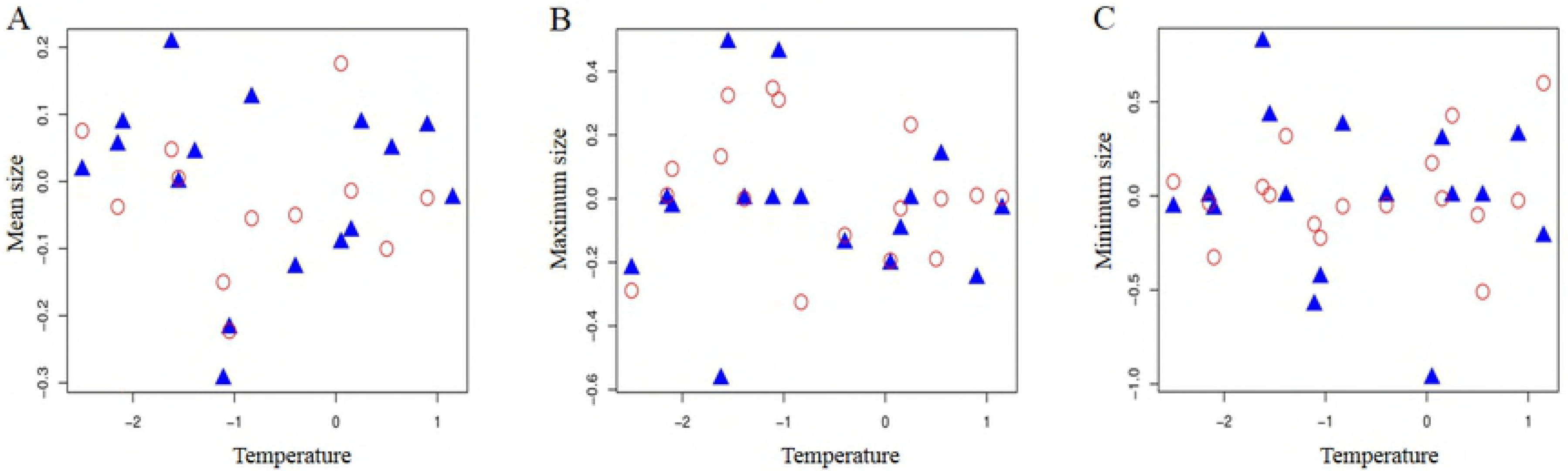
Climate induced body-size response has not been affected by the latitudinal spread. Dependence of climate induced body-size response on latitudinal spread of all (A), bivalves (B) or gastropods (C) families during Cenozoic Era. The vertical axis represents the coefficient associated with detrended correlation between temperature and mean body-size at family level. Size data is log transformed.

**Table 2.** Correlation ρ^1st^ _(Size-Temp)_ with existing body-size and latitudinal spread.

|  | Size dependence |  | Latitudinal dependence |  |
| --- | --- | --- | --- | --- |
| | $\rho^{1st}$ (Size- Temp) | P | $\rho^{1st}$ (Size- Temp) | P |
| Overall | 0.07 | 0.53 | 0.11 | 0.33 |
| Bivalve | 0.01 | 0.99 | -0.34 | 0.06 |
| Gastropod | 0.09 | 0.49 | -0.11 | 0.41 |
| Infauna | -0.20 | 0.38 | -0.51 | 0.01 |
| Epifauna | 0.13 | 0.30 | -0.12 | 0.36 |

The significance is set at a value determined by False discovery rate procedure.

There is no significant difference in the body size response to temperature between tropical and temperate species (Fig 5, Table 3).

**Fig 5.**
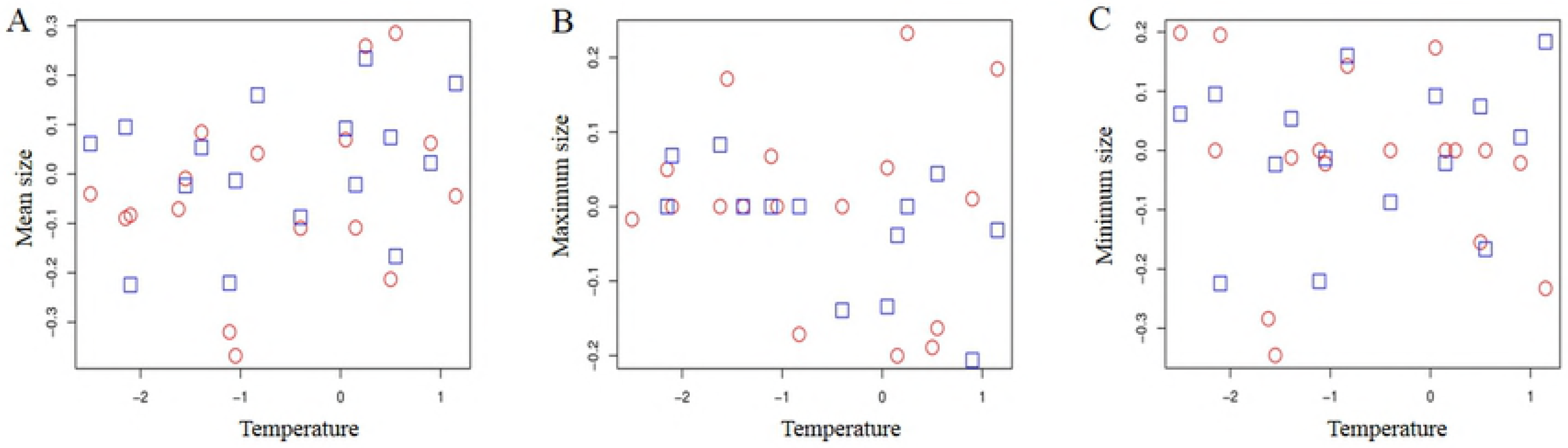
Detrended correlation between temperature and body-size do not show any difference in response between tropical and temperate species. The detrended relationship between body-size and temperature considering average body-size (A), maximum body size (B) and minimum body-size (C). The blue triangles represent tropical species and the red circles represent temperate species. Size data is log transformed.

**Table 3.** Results of Cramer test to evaluate the difference in response among species of varied latitudinal zones and ecology.

|  | Mean |  | Maximum |  | Minimum |  |
| --- | --- | --- | --- | --- | --- | --- |
| | $\rho^{1st}$ (Size-Temp) | P | $\rho^{1st}$ (Size-Temp) | P | $\rho^{1st}$ (Size-Temp) | P |
| Tropical- Temperate | 1.83 | 0.99 | 1.79 | 0.99 | 1.81 | 0.99 |
| Infaunal- Epifauna | 1.81 | 0.99 | 1.82 | 0.99 | 1.77 | 0.99 |

Both epifauna and infauna showed lack of correlation between temperature and body size in the detrended data. There is no significant difference in the body size response to temperature between infaunal and epifaunal species (Fig 6, Table 3). No correlation is observed between maximum body-size and ρ^1st^ _(size-temp)_ for either epifaunal or infaunal families (Fig 7A-B, Table 2). A negative correlation for infaunal families exists between ρ^1st^ _(size-temp)_ and latitudinal spread (ρ= −0.51; P= 0.01) (Fig 7C, Table 2); however, with false discovery rate procedure, this result is no longer significant implying infaunal families with limited latitudinal spread did not show any difference in the magnitude of body-size change compared to widespread taxa. Such latitudinal range dependence is absent among epifaunal families (ρ= −0.12; P= 0.36) (Fig 7D, Table 2).

**Fig 6.**
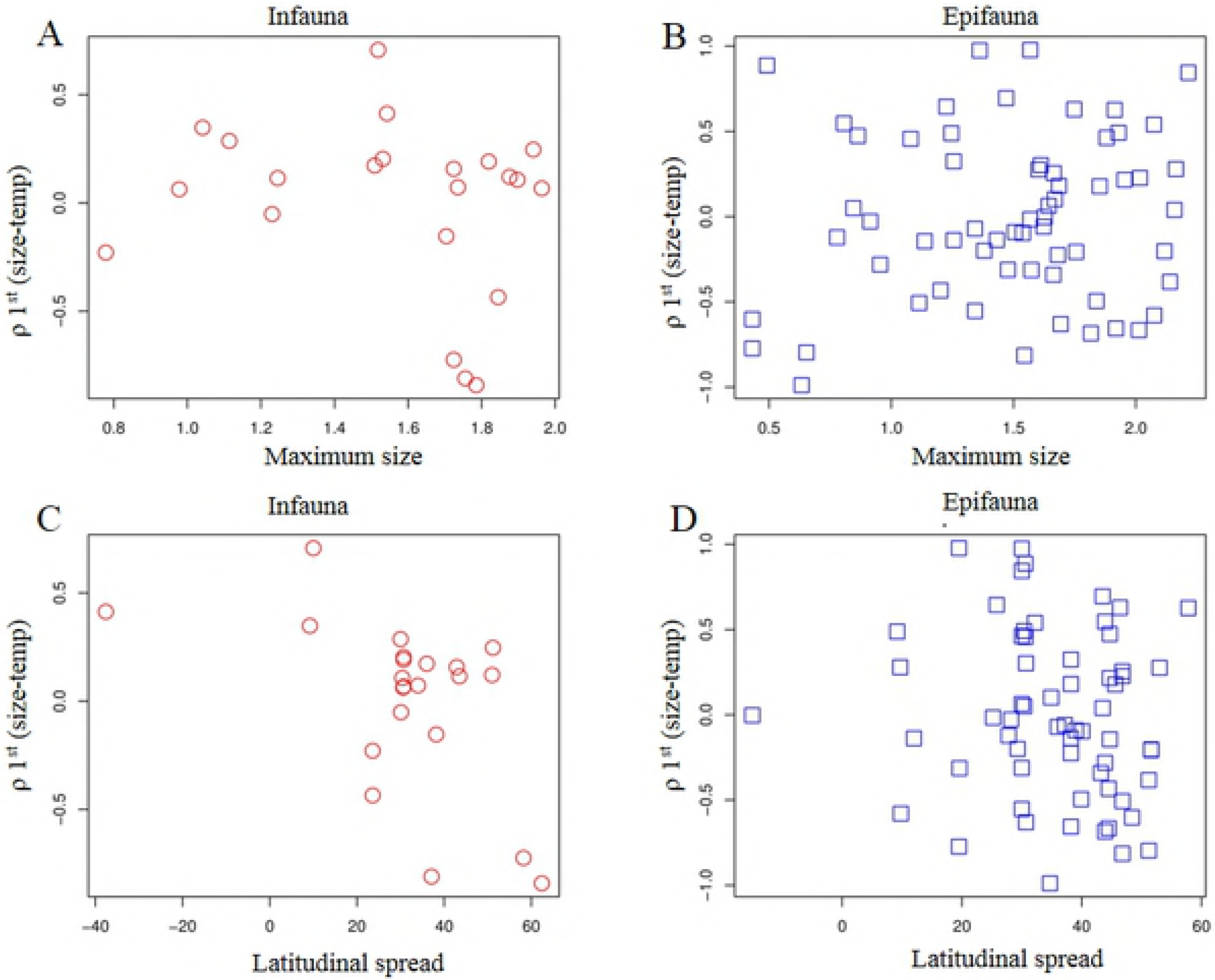
Detrended correlation between temperature and body-size do not show any difference in response between epifaunal and infaunal species. The detrended relationship between body-size and temperature considering average body-size (A), maximum body size (B) and minimum body-size (C). The red circles represent infaunal species and the blue squares represent epifaunal species. Size data is log transformed.

**Fig 7.**
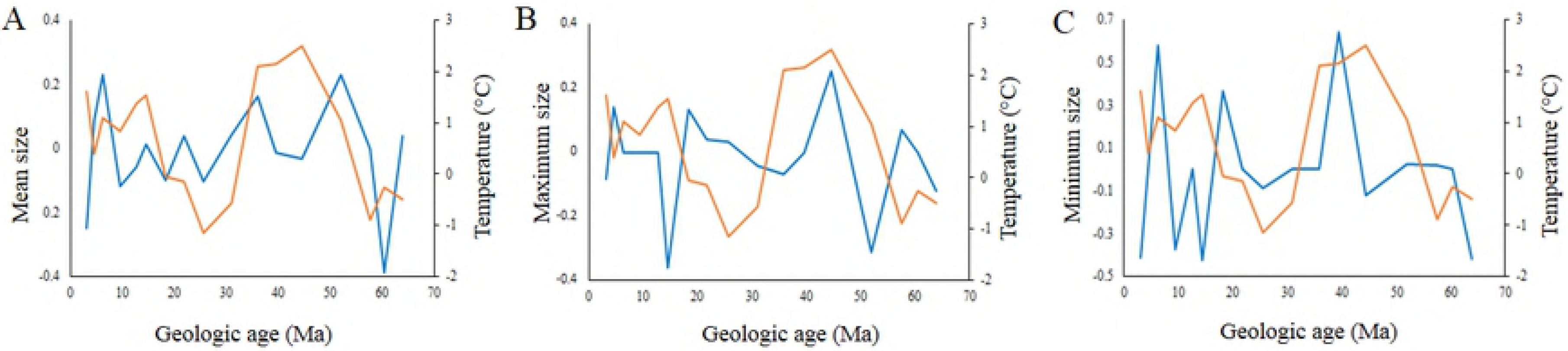
Climate induced body-size response has not been affected by the existing size or the latitudinal spread for infauna and epifauna. Dependence of climate induced body-size response on existing body-size of family for infauna (A) or epifauna (B). Dependence of climate induced body-size response on maximum latitudinal spread of family for infauna (C) or epifauna (D). The vertical axis represents the coefficient associated with detrended correlation between temperature and mean body-size at family level. Size data is log transformed.

## Discussion

In recently documented effect of Recent climate change on biota, reductions in body size has been suggested as the third universal response to global warming, along with changes in the phenology and distributions of species [17, 40–42]. A review of recent studies, however, also shows nonuniformity in the magnitude and direction of size responses across taxa and ecology [43]. This underscores the necessity to validate such claims with phylogenetically controlled comparative analyses of large temporal changes. Recent studies also point to a serious lack of comprehensive studies on marine ectotherms. Because, theoretical models of size-dependent thermoregulatory and metabolic responses are largely done on terrestrial endotherms, large scale analysis on body-size response of marine ectotherms is likely to contribute to our understanding of the underlying mechanisms of size shifts for such groups. This would, therefore, refine the ability to predict the sensitivities of species to climate change. Our study attempts to fill this gap by evaluating the body size response of marine fauna to climatic fluctuation in deep time.

### Absence of size reduction during events of climatic warming

The remarkable absence of the negative trend between body size and temperature in our observation of 66 My fossil record raises questions about the validity of TSR rule at species level for marine invertebrates. It is important to note, however, that the maximum size shows a negative relationship with temperature, albeit non-significant. There could be a number of reason for not finding the expected negative trend. The first possibility could be that the preservation bias may have destroyed the pattern that really existed [44]. However, the global nature of the database and relatively higher degree of preservation of this specific group, would reduce the chances of taphonomically induced changes in the result – a scenario that has already been established for the evolutionary patterns for this group [45]. More importantly, it would be hard to imagine a causal mechanism that would selectively remove a particular size class from a specific time bin, especially within a relatively narrow range of size spectrum (two orders of magnitude).

The second possibility is that the expectation of a species level TSR for marine molluscs is not valid. The idea of TSR at species level over a broad temporal span stems from widely accepted temperature–size relationships documented in geographically separated modern populations (Bergmann’s Rule). It has been argued that species and lineages that conform to Bergmann’s Rule should show a larger size during times of climatic cooling [10]. Marine molluscs have often shown latitude-dependent body size variation [46] and hence, expected to show the predicted negative relationship. The fact that they do not show the trend during Cenozoic climatic events probably indicates a more complex mechanism of body size evolution. Because body size is influenced by a variety of ecological and evolutionary tradeoffs between growth, resource availability, reproduction, predation, longevity among other factors, inter-specific patterns may fail to be simple extensions of intra-specific processes [19].

The third possibility could be related to the multifaceted nature of climate change itself and its complex influence on ecology. The mechanism of climate induced body size change may be influenced by other biotic and abiotic factors that may in turn have additive and multiplicative effects on size. A number of alternate studies have been proposed involving temperature-related variation in seasonality, growth constraints, oxygen requirement, and nutritional stress [47–50]. Hence, a lack of expected trend may arise because so many environmental factors are interacting during climatic shifts and not necessarily because of a lack of causal mechanism. Climate change, for example, is greatly associated with changes in nutrient availability apart from modifying temperature which may play a significant role in climate induced body size evolution during Cenozoic. For instance, Equatorial Pacific upwelling occurred in Eocene – a time of substantial climatic warming, increased the nutrient utilization and rate of metabolism [51]. Such intervals of higher nutrient availability would allow the development of larger body size [52] – a scenario which is in sharp contrast to the prediction of temperature-induced reduction in body size for the same time interval. On a similar line, Nawrot et al [53] demonstrated a size increase in the Mediterranean mollusc due to warming induced invasion. The effect of global warming indirectly led to an increase in the large bivalve species in the Mediterranean Sea by facilitating the entry and subsequent spread of tropical aliens. This study shows an outcome of increasing body-size during a time of warming that is opposite to the expectations based on the general TSR of marine ectotherms. Such contrasting outcomes from multiple events are likely to obscure the overall negative pattern of temperature-size relationship even if it exists at species level; hence, a complied dataset such as the present one may fail to show the expected negative relationship over broad temporal span.

Finally, the mechanism of body size change and speed of the process may play a role in dictating the nature of temperature-size relationship in deep time. A curve describing the relationship between a trait and the environmental variable - the reaction norm - has been predicted to have a negative slope for body-size and temperature. This slope depicts the sensitivity of the trait to the environment [54]. Such reaction norm is also characterized by an elevation i.e. the trait value in the average environment. A general temperature-size rule has been proposed to explain the pattern in ectotherms in the context of development reaction norms for size [55, 56], but the causes of size patterns remain hotly debated [57–60]. The controversy over size reductions stems from the debate on the nature of size shifts. Some argue that such size response represents evolved genetic responses to climate change and hence would be a slow process [61]. However, other researchers view it a phenotypic plasticity in response to changes in the surrounding environment (such as climate) and hence would be fast [62, 63]. This is an important issue because the likelihood that species can respond fast enough to climate change will ultimately depend on whether their response is genetic or plastic [42, 61, 64, 65]. Moreover, a phenotypically plastic response to temperature extremes would also mean a quick reversal of the trait value to the elevation of the reaction norm once the temperature changes to more average value. Such short lived morphological changes are less likely to leave a very distinct record in a time-averaged fossil succession.

### Lack of selectivity in the morphological response

Ohlberger et al [23] predicted that groups with narrow latitudinal spread should respond significantly to climatic fluctuation. However, we did not find any support for this claim. Lack of response could be due to a few issues. The first issue is related to the nature of our data and subsequent treatment. The family level data might be too coarse to document the pattern. The second issue could be related to the evolution of the thermal gradient through Cenozoic and its dissimilarity to modern one. The nature of thermal gradient between tropics and poles have changed over time. The strong gradient that we observe today, might have been absent in early parts of Cenozoic. Hence tropical fauna might not have behaved very strongly like today. The third issue could again be related to the type of response of a species to climatic fluctuation. A group can respond in different ways to a climatic fluctuation that includes change in the phenotype distribution via phenotypic plasticity [54], relocation to suitable habitats, and genetic change, i.e. microevolution [61, 66, 67]. The rate of adaptation without genetic change can be very high because phenotypic plasticity works in no time. If phenotypic plasticity is working alongside relocation of groups to suitable habitats, the resulting changes should not be expected to happen at any specific locale.

The relative increase in metabolic costs with temperature is greater in warmer climates such as tropics due to the exponential increase in metabolic rate, and thermal windows tend to be narrower in tropical compared to temperate environments due to lower temperature variability [68, 69]. Tropical species, therefore, typically have an upper thermal limit for survival closer to the optimum temperature than their temperate counterparts. Temperate species have broader thermal tolerances and generally experience climates with average temperatures below their thermal optima [68, 70]. Based on this, Ohlberger et al [23] predicted that groups with tropical affinity should respond significantly to climatic fluctuation. Contrary to this prediction, our data shows that the body-size response of temperate species is similar to the response of tropical species. Such lack of tropical sensitivity in response can be explained by other factors controlling body-size. Another cause of changing body size apart from temperature is the change in the availability or quality of food that, in turn, affects nutrition, and this has been implicated as a mechanism in the majority of the studies to date [61]. Changes in nutrition could result from changing temperature, for example via changes in the length of growing season [71, 72] or via changes in temperature-dependent activity budgets that constrain feeding [73] in terrestrial habitats. In the ocean, however, nutrient distribution and productivity is controlled by a combination of global and regional factors. Moreover, specific marine provinces are characterized by different combinations of temperature and productivity [74, 75]. Hence, biotic response of the shallow marine fauna should be strongly dependent on regional variation of annual SSTs [75], pointing to the role of provincial nature of the marine thermal structure in shaping the productivity profiles. The combined effects of the seasonality of solar energy outside the tropics, increasing with latitude, and the episodic nature of upwelling at any latitude, creates a positive correlation between seasonal variation and mean annual values in productivity, but one that is not linearly related to latitude [76]. In spite of such regional variations, solar radiation is the primary controlling factor in making the surface water warmer and less dense than deeper waters. Such stratification is highest in the tropical shelves with the greatest density difference between surface and deeper waters. This stratification promotes stability within the shallow water column, inhibiting vertical water motion and tending to stabilize productivity in the euphotic zone [77]. The shallow tropical waters, thermally stratified and thus relatively stable, are often relatively nutrient poor, and productivity is therefore low but also tends to be relatively stable. Consequently, low latitudes are characterized by highest SST in the shallow sea, but not the productivity; higher mid-latitudes show highest productivity values. Because of the relative stability in productivity in tropics, tropical fauna is less likely to show severe body-size response, unlike the prediction based solely on temperature induced morphological changes. The major influence of regional character of ocean basin and the interaction of those factors is likely to be important in guiding the body size trajectory of marine species than simplistic general rules predicting heightened response of tropical species.

### Absence of ecological selectivity

The ecological selectivity of recent global warming is one of the most debatable issues. The complex interplay of ecology and climate change in shaping macroevolutionary pattern of marine groups has been documented for Cenozoic [78] highlighting the importance of ecological attribute of a group in dictating various physiological response to abiotic factors. There have been predictions about differential morphological response of various ecological groups of molluscs due to their habitat. Thorson [29] claimed that infaunals are shielded from temperature fluctuation because of burrowing and hence they would not show the classical LBG. According to this postulation, the infaunals are less likely to show any body size response to climatic variation due to such shielding in comparison to epifaunals that are always exposed to temperature change. However, an opposite argument can be made from the point of inherent difference in the thermal specialization [26]. Exposure to very high or low temperature for a short period usually causes increased tolerance to acute thermal extremes whereas long-term exposure to moderate temperatures can induce heightened thermal sensitivity [25, 26]. This mechanism has been used to explain the observed difference in body size reduction between marine and terrestrial groups as a response to temperature. Most terrestrial ectotherms experience stronger short-term variability than marine species due to their lower thermal buffering and show a less pronounced reduction in body-size in response to warming compared to those of the marine organisms [79]. Similarly, epifauna facing larger variation in temperature, should show a higher temperature tolerance and less sensitivity to climate induced body-size change unlike infaunas that are acclimatized to a narrow temperature zone and likely to show a greater response due to their higher thermal sensitivity [14]. This difference should be more pronounced for infauna of narrow latitudinal extent - both of these factors are known to generate lower thermal tolerance [14]. Our results, however, does not demonstrate strong ecological selectivity in controlling the sensitivity of climate-induced morphological change pointing to limitation of these generalized predictions.

## Conclusions

We showed that generalized body size reduction to climatic warming may not have existed at species level for marine benthic molluscs during Cenozoic events of climatic fluctuation. The prediction of heightened response of the groups with narrow latitudinal range, tropical affinity or with higher body size, also fails to show consistent pattern in their body-size response during the past 66My long climatic fluctuation. In spite of the general predictions of higher response of tropical species, our data do not demonstrate any significant difference in body-size response between tropical and temperate species during the climatic fluctuations of Cenozoic. Based on the prediction based on thermal specialization, molluscs with varying substrate relationship are expected to respond differently with climatic change. Our comparison between epifaunal and infaunal molluscan group, however, does not show any support for this prediction. The two groups do not show significant difference in their body-size response during times of climatic fluctuation of the past. Although, infaunal families with narrow latitudinal spread showed higher degree of climate influenced size variation compared to the epifaunals, this pattern in not statistically significant. These observations are pointing to a complex interplay between temperature and other regional factors during climatic fluctuation and the limited validity of “universal rules” in governing the species response in marine ecosystems. These results underscore the complex nature of climate induced morphological change among marine species and the need for detailed regional study to evaluate the controlling factors in the absence of a universal rule governing marine ecosystems.

## Acknowledgements

This project was supported by Academic Research Grant, IISER Kolkata (ARF 2015-16) and IISER Kolkata graduate research fellowship. Comments from two anonymous reviewers on a previous draft greatly improved the manuscript. We would like to thank Satyaki Mazumdar for his help in statistical analysis.

## Supplementary materials

### S1 Data. Dataset used for the present study

Collection_no: Number assigned to each specimen in PaleoDB, Species: Species name, Family: Family name, Class: Class name, Specimens_measured: Number of specimens used to take measurements, Measured value: Geometric mean of the measured specimens available in PaleoDB, Size (log transformed): Log transformed value of “Measured value”, Latitude: Paleo-latitude of the area from where the specimen was collected. A negative value indicates southern hemisphere, U_time_int: Upper limit of the time interval, Time bin: Geologic time intervals based on age unit, Habitat: Substrate relationship of each species, Zone: Latitudinal zone of occurrence of a species, Temperature: Average temperature of each time bin.

**S2. R script for statistical analyses used for the present study**

